# Graph-based RNA structural representation reveals determinants of subcellular localization

**DOI:** 10.64898/2026.02.23.707397

**Authors:** Yi Hao, Heyun Sun, Zixu Ran, Xudong Guo, Ming Liu, Yue Bi, Jose M Polo, Ning Liu, Fuyi Li

## Abstract

RNA subcellular localization is a key determinant of RNA function and regulation, yet existing computational approaches rely primarily on sequence or simplified structural descriptors, limiting their scalability to long transcripts, their ability to model inter-label dependencies, and their applicability across RNA types. Here, we present GRASP, a unified graph neural network framework for predicting RNA subcellular localization using a heterogeneous graph representation that is RNA substructure-aware. GRASP presents each RNA as a multi-scale graph comprising nucleotide nodes and secondary-structure-derived substructure nodes, connected by relational edges, enabling joint modeling of base-level interactions and regional structural context. The model further incorporates multi-label dependency learning to capture co-localization patterns across cellular compartments within a unified framework. Across multiple benchmark datasets and RNA types, GRASP consistently outperforms state-of-the-art sequence-based and structure-informed methods, achieving substantial improvements in accuracy, F1 score, and AUC while maintaining strong scalability to long transcripts. In addition, the graph-based representation provides biologically interpretable insights into structural determinants of RNA localization. The source code and data are available at https://github.com/ABILiLab/GRASP, and the web server is accessible at http://grasp.biotools.bio.

## 1 Introduction

RNA molecules, including messenger RNAs (mRNAs) and long non-coding RNAs (lncRNAs), play essential roles in gene and cellular regulation, as well as in disease processes (1,2). Increasing evidence indicates that RNA function is tightly coupled to its subcellular localization. For instance, mRNA localization influences translational efficiency and protein targeting (3,4), whereas lncRNA localization determines regulatory activity in chromatin remodeling, transcriptional, and post-transcriptional control (5,6). More broadly, localization constrains which binding partners and regulatory machineries an RNA can access, thereby shaping core steps of RNA metabolism, including splicing and nuclear export, translation, and RNA turnover. Experimental approaches such as fluorescence in situ hybridization (FISH) (7) and fractionation-sequencing (8) can directly measure RNA localization, but they remain costly, labor-intensive, and often condition-dependent, making comprehensive profiling difficult to scale to the rapidly expanding catalogue of newly discovered transcripts. These limitations motivate the development of computational methods for accurate, transcriptome-wide prediction of RNA subcellular localization.

Early computational methods primarily relied on engineered sequence features combined with classical machine learning classifiers. Representative examples include mRNALoc (3) and LncLocPred (9), which employ hand-crafted descriptors and conventional models to predict localization. With the rise of deep learning, neural network-based approaches began to automatically learn sequence representations, such as DeepLncLoc (10) and iLoc-lncRNA (11). More recently, transformer architectures have been introduced to capture long-range dependencies in RNA sequences, such as LncLocFormer (13), and several studies, including our Allocator (16) and LncTracker (17), have attempted to incorporate RNA structural information using graph neural networks or hybrid multi-modal frameworks (14,15). These advances have also shifted prediction settings from single-label classification to more realistic multi-label formulations (12).

Despite these advances, several key challenges remain. First, most existing methods still rely predominantly on linear sequence representations, even though RNA secondary structure plays a critical role in localization. Structural information, when included, is typically incorporated as auxiliary descriptors rather than being modelled as a primary representation, limiting the ability to capture higher-order structural context and hindering scalability to long transcripts. Second, localization labels are commonly treated as independent prediction tasks, ignoring biologically meaningful dependencies among cellular compartments. Third, many existing tools are specialized for a single RNA class, restricting their generalizability across RNA types.

To address these limitations, we present GRASP (**G**raph-based **R**NA Substructure-**A**ware **S**ubcellular localization **P**rediction), a unified graph neural network framework for RNA localization prediction. GRASP represents RNA molecules as heterogeneous graphs composed of nucleotide nodes and secondary-structure-derived substructure nodes connected by relational edges, enabling joint modelling of base-level interactions and region-level structural context within a single architecture. The framework further incorporates label-dependency modelling to capture co-localization patterns across cellular compartments. Extensive experiments demonstrate that GRASP improves predictive accuracy, interpretability, and generalizability across both mRNA and lncRNA localization tasks.

## 2. Materials and methods

### 2.1 Overview of the Framework

The GRASP framework is designed as an end-to-end pipeline for multi-label RNA subcellular localization prediction (**Figure 1**). The overall workflow consists of three major stages: dataset construction, multi-modal feature representation, and multi-branch model learning. First, RNA sequences and localization annotations are curated to construct benchmark datasets covering both mRNAs and lncRNAs. Each RNA sequence is then transformed into a substructure-aware heterogeneous graph, which explicitly represents nucleotides together with secondary-structure-derived substructures. In parallel, complementary sequence-level features are extracted to capture compositional and long-range dependency information. Second, GRASP adopts a multi-branch architecture to process heterogeneous feature modalities. The heterogeneous RNA graph is encoded using a graph neural network to learn structure-aware representations, while sequence-derived features are processed through dedicated feed-forward channels. The resulting embeddings are integrated to produce a unified representation for each RNA molecule. Finally, the model is trained under a multi-label learning framework that explicitly incorporates label correlations through a regularization term and addresses class imbalance using asymmetric loss. This design enables GRASP to jointly model RNA structural context, sequence characteristics, and inter-label dependencies within a unified prediction framework.

**Figure 1.**
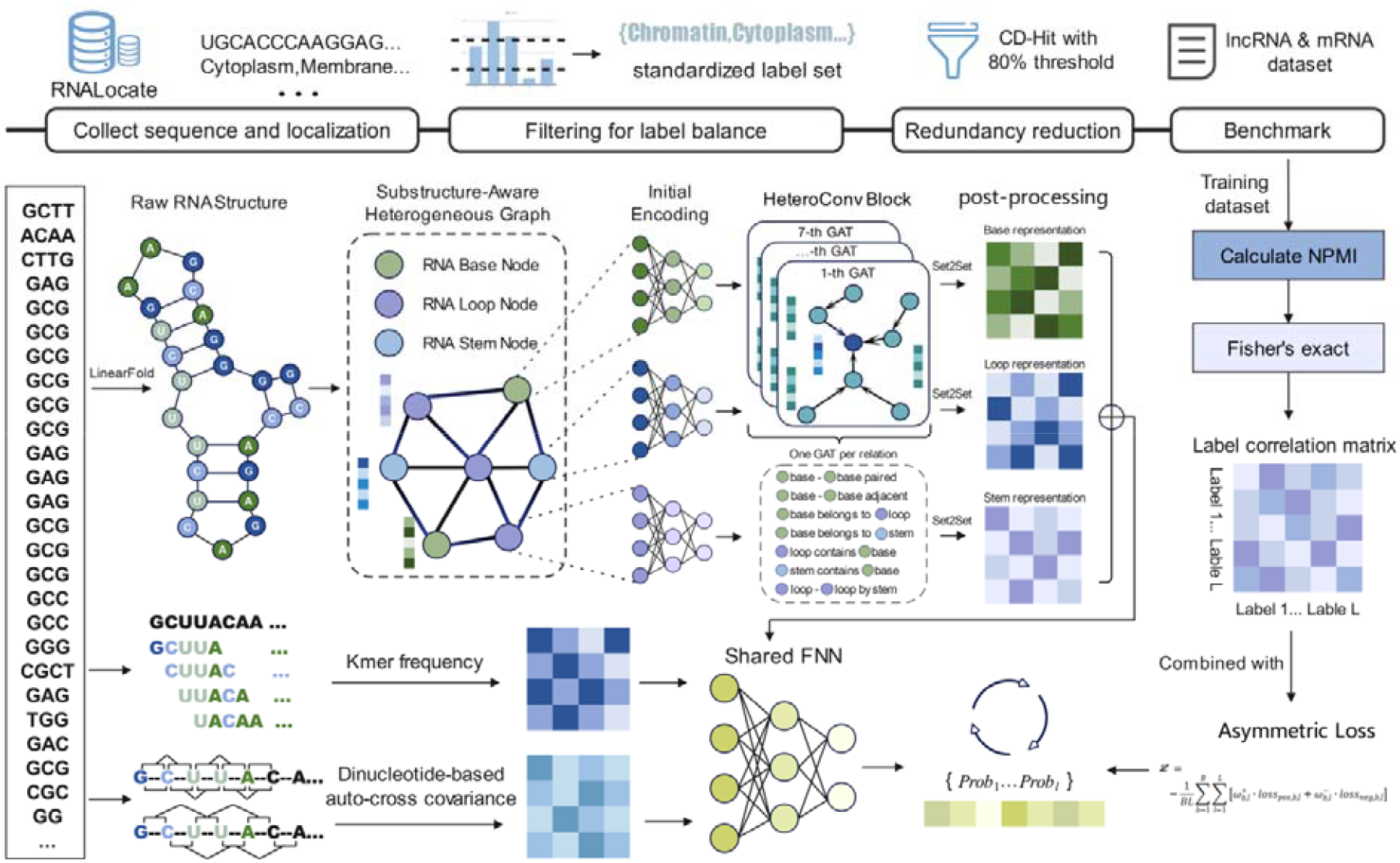
Overview of the GRASP framework. The workflow includes dataset construction, substructure-aware heterogeneous graph representation of RNA, extraction of complementary sequence features, multi-branch neural network integration, and training with an asymmetric loss that incorporates label-correlation constraints for multi-label localization prediction.

### 2.2 Dataset construction

To develop and rigorously evaluate GRASP, we construct benchmark datasets for RNA subcellular localization by curating experimentally validated data from publicly available resources and previously published studies. Both messenger RNAs (mRNAs) and long non-coding RNAs (lncRNAs) are included to ensure broad applicability across major RNA classes. RNA sequences along with their curated subcellular localization annotations across multiple cellular compartments are collected from the RNALocate 3.0 database (18). For RNAs with multiple reported localization records, annotations are merged into a unified multi-label format and matched to their corresponding RNA sequences. This initial collection comprised 12,118 lncRNA entries and 212,855 mRNA entries. To mitigate severe label imbalance in the multi-label setting while preserving a realistic class imbalance, we select localization categories that are either extremely rare or overly dominant, together with sequences exclusively associated with those labels. To further reduce sequence redundancy, we use CD-HIT (19) with an 80% sequence identity threshold, ensuring that the datasets remained representative while avoiding overrepresentation of highly similar sequences. After preprocessing, the final lncRNA dataset (LncDS) contains 6,654 sequences annotated with eight subcellular localizations: cytoplasm, cytosol, extracellular vesicle, membrane, mitochondrion, nucleolus, nucleoplasm, and ribosome. The final mRNA dataset (mDS) contains 30,912 sequences annotated with eight compartments: chromatin, cytoplasm, cytosol subcompartment, membrane, microvesicle, nucleolus, nucleoplasm, and nucleus subcompartment. These curated datasets form the basis for all subsequent model training and evaluation.

### 2.3 Construction of the substructure-aware heterogeneous graph

To systematically capture both RNA sequence and secondary-structure context, each RNA molecule is represented as a substructure-aware heterogeneous graph. In this graph, three node types are defined: nucleotide bases (base), loop regions (loop), and stem regions (stem). Edge types are designed to represent nucleotide adjacency, base-pairing interactions, hierarchical membership relations, and connectivity between different structural elements. The construction scheme of the heterogeneous graph is illustrated in **Figure 2**. We employ dot-bracket notations of the RNA secondary structures predicted by LinearFold (20) in this study. Base types and pairing states can be naturally derived from this representation. To capture loop and stem substructures, we propose a recursive algorithm (**Algorithm 1**) that scans the dot-bracket string and iteratively extracts nested loop-stem units. The algorithm begins by identifying the outermost unpaired regions to form loop nodes, then detecting contiguous base-paired segments that define stem nodes. Nested loops are recursively parsed until the full structure is decomposed into hierarchical substructures. This procedure yields a structured set of bases, loops, and stems that serve as the foundation for heterogeneous graph construction.

**Figure 2.**
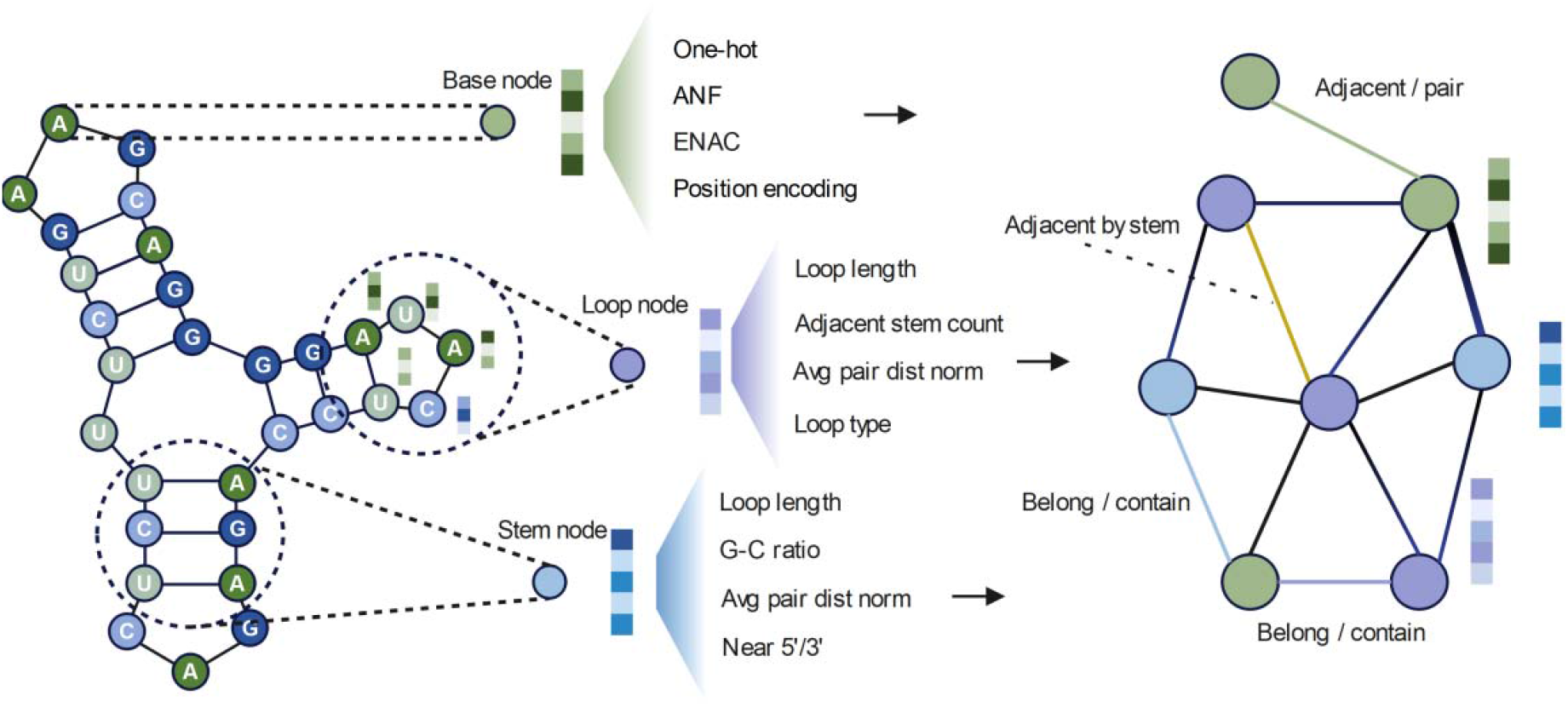
Schematic illustration of the substructure-aware heterogeneous graph construction. The diagram highlights the three node types (base, loop, and stem), their sources and feature representations, and the diverse edge types used to encode sequence adjacency, base-pairing interactions, and structural relationships among RNA substructures.

Specifically, after obtaining RNA secondary structures, we construct substructure-aware heterogeneous graphs. Formally, each sequence is transformed into a heterogeneous graph defined as:

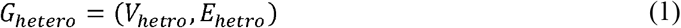

where *V*_*hetro*_ and *E*_*hetro*_ denote the sets of nodes and edges in the heterogeneous graph, respectively.

#### Algorithm 1

Construction of Substructure-Aware RNA Heterogeneous Graph

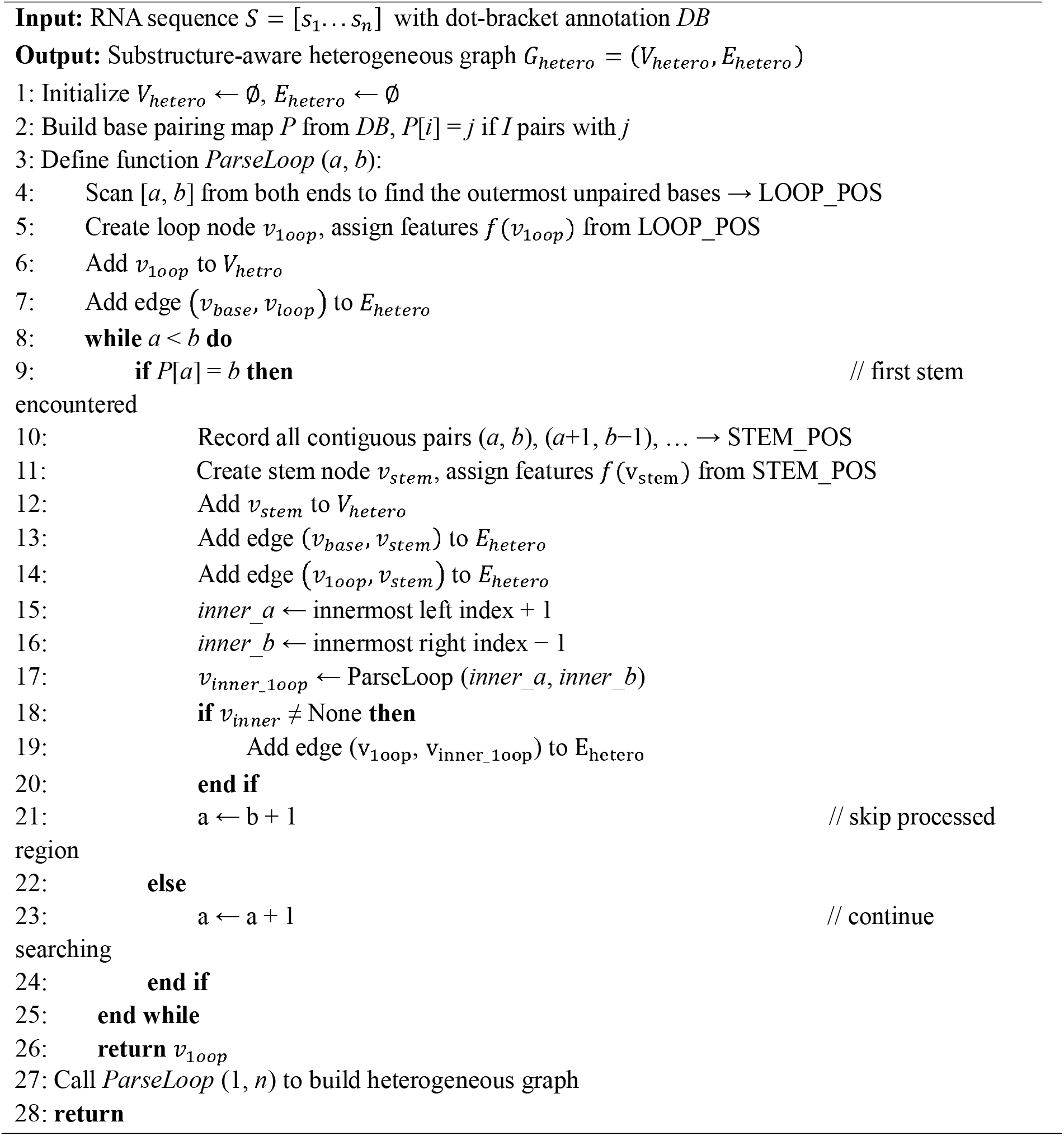

The node set consists of three types of nodes, including,, and, representing different hierarchical structural units. To encode prior information of each node type, we assign distinct initial features. The initial features of base nodes include one-hot encoding of nucleotide type, accumulated nucleotide frequency (ANF), enhanced nucleic acid composition (ENAC), and sinusoidal positional encoding, which can be expressed as:

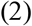

The initial features of loop nodes include the normalized number of bases in the loop, the normalized number of adjacent stems, the normalized average base-pair distance within the loop, and a one-hot encoding of loop types (internal loop, multi-loop, and hairpin loop), represented as:

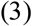

The initial features of stem nodes include the normalized number of bases *n*_*stem_base*_, G-C pairs content *n*_*gc*_, normalized average base-pair distance within the stem *n*_*stem_pair*_, and indicators for proximity to the 5’ or 3’ ends, expressed as:

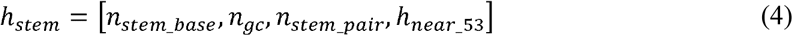

The edge set comprises seven types of edges to model the complex interactions in the secondary structure:

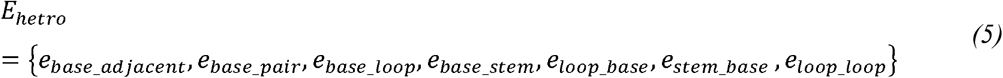

where *e*_*base_adjacent*_ represents adjacency between bases, *e*_*base_pair*_ represents complementary base-pair interactions, *e*_*base_loop*_ and *e*_*base_stem*_ denote directed membership relations from base nodes to loop or stem nodes, *e*_*loop_base*_ and *e*_*stem_base*_ represent the corresponding directed containment relations from loop or stem nodes to their constituent bases, and *e*_*loop_loop*_ represents loop-to-loop connections mediated by stems. Through these nodes and edges, the heterogeneous graph can simultaneously encode both the sequence information and the relationships among secondary structure subunits of RNA, providing a comprehensive structural context for subsequent graph neural network processing.

### 2.4 Complementary sequence-derived feature representation

In addition to the substructure-aware heterogeneous graph representation, the proposed framework incorporates two complementary sequence-derived features: *k*-mer composition and dinucleotide-based auto-cross covariance (DACC), to capture intrinsic sequence patterns.

The *k* -mer representation encodes the normalized frequency of contiguous subsequences and reflects local compositional characteristics of RNA sequences. For an RNA sequence *S*= [*s*_1_,*s*_2,_,… *s*_*n*_] the *k* -mer feature for pattern *u* is defined as:

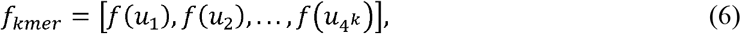

where

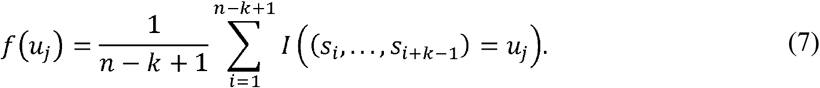

Here, *n* denotes sequence length, *k* is the *k* -mer size, *u* represents a specific *k* -length nucleotide pattern, and *I*(·) is the indicator function.

To further capture long-range sequence-order dependencies, we employed DACC features to characterize correlations between dinucleotide physicochemical properties across different positional lags. The DACC value is computed as:

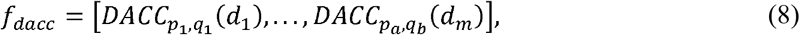

where each component corresponds to the covariance between dinucleotide physicochemical properties *p*_*a*_ and *p*_*b*_ at a predefined positional lag *d*_*m*_. Each element is computed as:

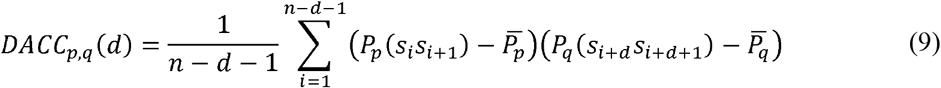

Where *p*_*p*_ (·) and *p* _*q*_(·) represent the values of physicochemical properties *p* and *q* for a given dinucleotide, while 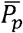 and 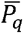 are the corresponding mean values over the entire sequence. All sequence features were extracted using the *iLearn* toolkit (21).

### 2.5 Modelling label dependencies and loss regularization

RNA subcellular localization prediction is inherently a multi-label task in which localization categories frequently co-occur or exhibit mutual exclusivity. To explicitly model these dependencies, we construct a label correlation matrix from the training labels *Y* ∈ {0,1} ^*N*×*L*^and incorporate it as a regularization term in the loss function.

The correlation between label pairs (*i, j*) is quantified using pointwise mutual information (PMI) and its normalized form (NPMI):

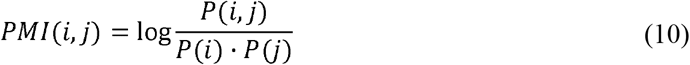

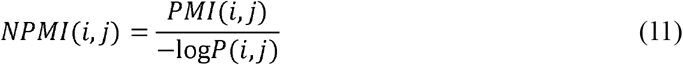

where *P* (*i*)and *P (i,j)* are estimated from the empirical label distribution *Y*. To reduce noise and eliminate spurious associations, Fisher’s exact test (22) is applied and only the top-*k* strongest correlations per label are retained. This procedure produced a sparse correlation matrix *R* ∈ [−1,1]^*L*×*L*^, encoding both positive co-occurrence (positive) and mutual exclusivity (negative). To incorporate these dependencies during training, we introduce a correlation regularization term that constrains predicted label probabilities:

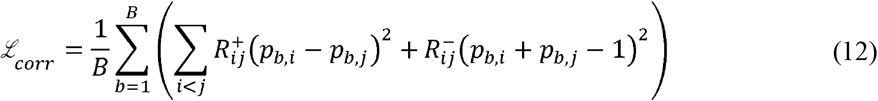

where *p*_*b,i*_ is the predicted probability for label in sample *b*, 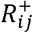 and 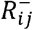 denote the positive and negative parts of *R*, and *B* is the batch size. Notably, we sum only over upper-triangular entries. Intuitively, this loss encourages positively correlated labels to have similar predicted probabilities and discourages the simultaneous activation of negatively correlated labels.

To address label imbalance in this multi-label localization data, we adopt Asymmetric Loss (ASL) (23):

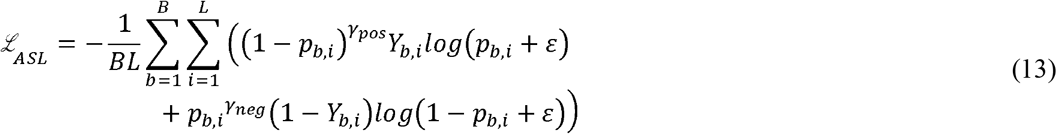

where *Y*_*b,i*_ is the true value for label in sample *b, γ* _*pos*_ and *γ*_*neg*_ control the focusing of positive and negative samples separately, ε is a numerical stability term. The final objective combines ASL with correlation regularization:

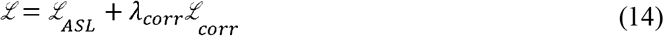

where *λ*_*corr*_ is the regularization weight. This design enforces label correlations while addressing imbalance, improving both prediction accuracy and reliability.

### 2.6 Model design

To effectively integrate the heterogeneous graph representation with sequence-derived features, we designed a multi-branch architecture that processes each modality through dedicated channels before feature fusion. Each node type in the heterogeneous graph, including base, loop, and stem, has its own initial feature vector, whose dimension differs and is relatively low. To unify the feature dimension, we first applied a type-specific multi-layer perceptron (MLP)(24) to project each node’s raw feature into a common hidden dimension *d*, In addition, to enhance model flexibility and allow the network to learn beyond handcrafted features, we concatenate a learnable embedding vector of the same dimension with the encoded feature. The initial encoding process can be formulated as:

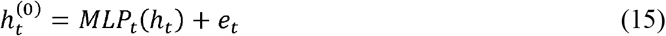

where *h*_*t*_ is the initial feature of the node of type *t* ∈ {*base, loop, stem* }, *MLP*_*t*_ (·) denotes the node-type-specific encoder, and *e*_*t*_ is a learnable embedding vector of the same dimension.

We then applied a heterogeneous graph convolution network based on the Graph Attention Network (GAT) (25) principle. In this setting, heterogeneous message passing is performed by learning relation-specific attention mechanisms, allowing the model to selectively weight information from different types of neighboring nodes. For each edge type *r* ∈ *R*, a dedicated attention-based operator *GAT*_*r*_ was employed to aggregate messages from neighboring nodes:

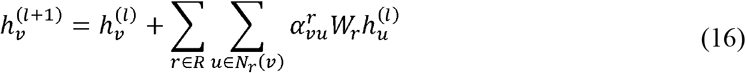

where *N*_*r*_ ( ν) denotes the neighbors of node *ν* under relation *r, W*_*r*_ is a learnable linear transformation matrix specific to relation *r*, and 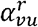 is the attention weight computed as:

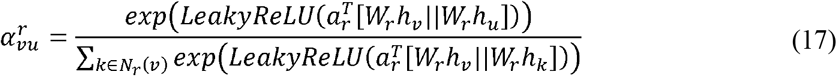

where *ν* and *u* denote the target node and one of its neighboring nodes under relation type *r*, respectively, h_v_ and h_u_ are the input feature vectors of nodes *ν* and *u, a*_*r*_ is a relation-specific attention weight vector, and ||indicates concatenation. After *L* layers of heterogeneous message passing, node embeddings of each type are pooled via Set2Set(26) to obtain a fixed-size graph representation:

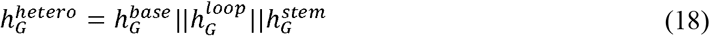

Where 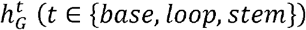 denotes the pooled vector for node type.

The two sequence-derived features, *k*-mer and DACC, are processed independently using simple yet efficient feed-forward networks:

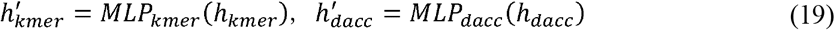

where *h*_*kmer*_ and *h*_*dacc*_ are the vectorized representations of the *k*-mer and DACC features, respectively.

Finally, the graph-level embedding and the two sequence-level embeddings are concatenated and fed into a shared feed-forward network to obtain the final prediction vector:

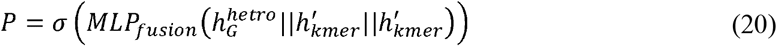

where *P* represents the predicted probabilities for all L labels, and *σ* is the element-wise sigmoid function.

### 2.7 Training strategy and evaluation metrics

We adopted a five-fold cross-validation protocol to ensure reliable evaluation. The model was trained with a batch size of 64, an initial learning rate of 0.001, and up to 200 epochs. An early stopping strategy was applied, terminating training if the validation loss failed to improve for 20 epochs. A cosine annealing scheduler with warm restarts was used to adjust the learning rate dynamically. The loss function combined ASL with a label-correlation regularization term, with a weight of 0.3 to prevent over-regularization. To achieve a more comprehensive evaluation, we adopted example-based and label-based metrics. The example-based metrics include example accuracy, Hamming loss, zero-one loss, coverage error, ranking loss, and average precision score; the label-based metrics include Matthew’s correlation coefficient, average F1-score, micro precision, micro recall, and average AUC. Detailed definitions and calculation procedures of all evaluation metrics are provided in the **Supplementary Materials**. All experiments were conducted on a workstation equipped with a single NVIDIA RTX 4090 GPU (24 GB memory) and a 12-core CPU.

## 3. Results

### 3.1 Benchmarking study

To evaluate GRASP’s performance in RNA subcellular localization prediction, we conduct an extensive benchmarking study against established methods, including representative machine- and deep-learning approaches. Since most prior studies focused on a single RNA type (mRNA or lncRNA) and thus cannot be directly compared with GRASP, we perform a stratified comparison by RNA type and the corresponding specialized studies. For the mRNA subcellular localization prediction task, mRNALoc (3), DM3Loc (27), Allocator (16) are selected as benchmark methods, while for the lncRNA prediction task, LncLocator2.0 (28), DeepLncLoc (10), LncLocFormer (13), and LncTracker (17) are used for comparison. Notably, for a broad comparison, some multi-class prediction tools, such as mRNALoc, LncLocator2.0, and DeepLncLoc, are also included, and we standardize all tools to use 8-class output heads with a Sigmoid activation function. The results of the benchmarking study are shown in **Figure 3a**.

**Figure 3.**
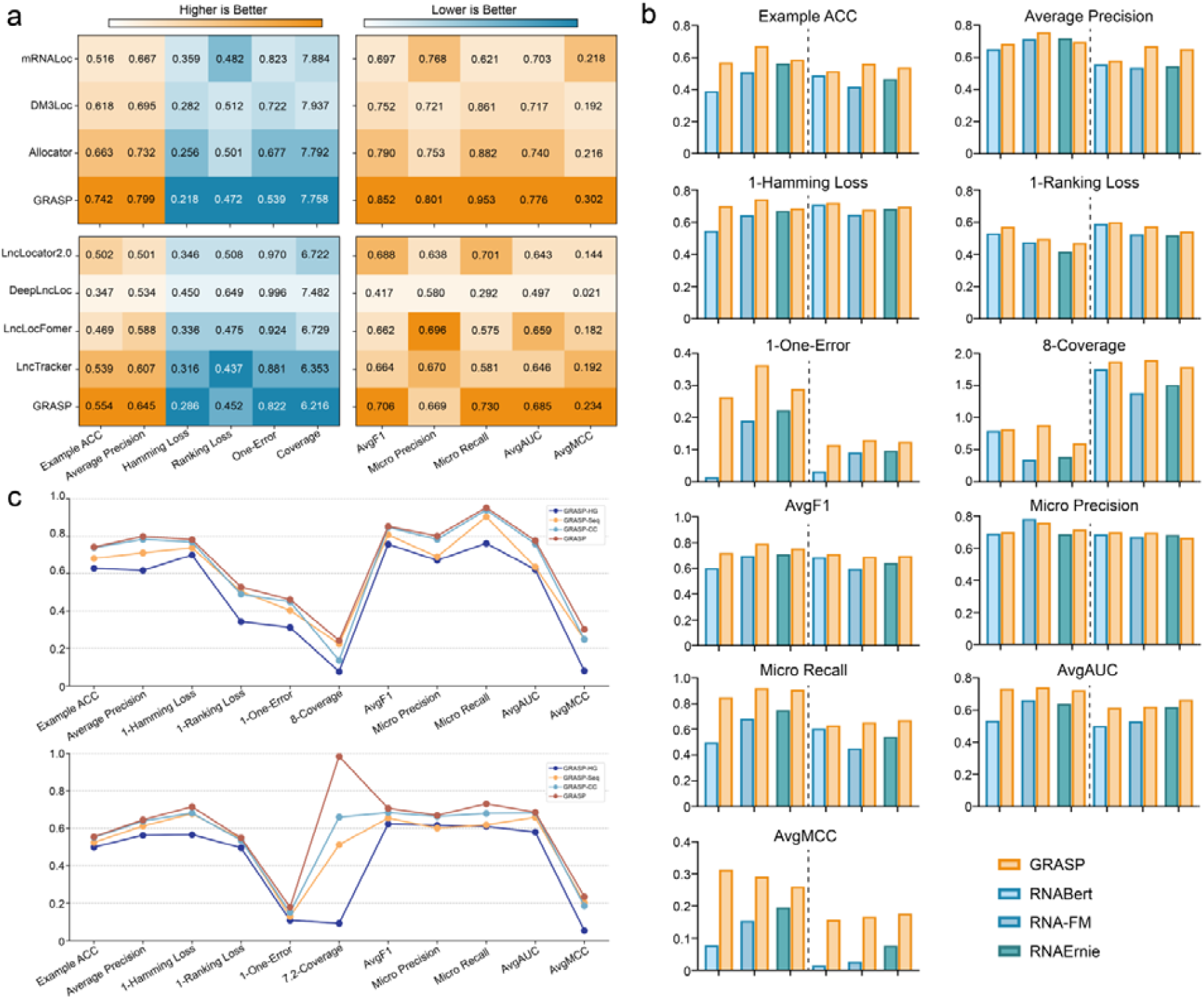
(a) Comparison of GRASP with representative subcellular localization models on the benchmark dataset. (b) Comparison of GRASP with existing RNA foundation models on a length-limited dataset, each subfigure is split into two parts, left for mRNA, right for lncRNA. (c) Ablation study results showing the performance of different GRASP model variants, highlighting the contribution of each module.

On the mRNA dataset, GRASP outperforms other methods in all example-based metrics, with the most notable improvement in One-error (0.539), which surpasses the second-best method, Allocator (0.677), by 0.138 (Figure 3.a). GRASP also reduces Hamming loss (0.218 vs. 0.256) and achieves a lower Ranking loss (0.472 vs. 0.482), indicating fewer overall prediction errors and more accurate label ranking for individual samples. GRASP also consistently achieves the best performance in all label-based metrics compared to other methods. The most significant gain is observed in AvgMCC (0.302 vs. 0.218), underscoring GRASP’s robustness and generalization in multi-label classification. On the lncRNA dataset, GRASP consistently achieves the highest average precision (0.645), clearly exceeding the second-best method (LncTracker, 0.607). In the label-based evaluation, the most significant improvement remains in AvgMCC (0.234), which is a clear improvement over the best baseline (LncTracker, 0.192). Collectively, these results demonstrate GRASP’s superior performance in multi-label prediction of subcellular localization across both mRNA and lncRNA datasets.

### 3.2 Comparison with RNA foundation model

To further evaluate the generalization ability of GRASP, we compare its performance with several representative RNA foundation models, including RNABERT (29), RNA-FM (30), and RNAErnie (31). For downstream classification evaluation of these pretrained language models, a unified fine-tuning strategy is adopted, sequence representations are extracted using the designated pretrained models and feed into a lightweight classification head for prediction. During fine-tuning, to assess their task adaptation capability under limited supervised signals, the backbone parameters of the language models are frozen, and only the classification head is updated. Given that different RNA foundation models impose distinct constraints on input sequence length (440 nt for RNABERT, 1022 nt for RNA-FM, and 512 nt for RNAErnie), we construct model-specific length-restricted datasets for a fair comparison. Specifically, sequences whose lengths did not exceed the maximum input limit of each model are selected from the original benchmark dataset, and performance comparisons are conducted exclusively on the corresponding restricted test sets. For the RNA foundation models, 60% of the sequences in each length-restricted dataset are used to fine-tune the classification head, while the remaining 40% are reserved for testing. For GRASP specifically, we keep the test set identical to that used for the foundation models, use remaining sequences in the dataset for model training, therefore fully leveraging its task-specific modeling capacity and structural information integration. As a result, with specialized task settings, we observe that GRASP consistently exhibits more stable and competitive performance than general-purpose RNA foundation models (**Figure 3b**), highlighting its advantages of task-driven modeling across different dataset settings.

### 3.3 Ablation study

To evaluate the contribution of each component in GRASP, we conduct a series of ablation experiments by systematically removing or modifying specific modules. All model variants are trained and evaluated on the same mRNA and lncRNA datasets using both example-based and label-based metrics to ensure a comprehensive assessment of predictive performance. The evaluated variants include: *GRASP*_*-HG*_ (without heterogeneous graph features), *GRASP*_*-Seq*_ (without sequence features), and *GRASP*_*-cc*_ (without the label correlation constraint). The results are summarized in **Figure 3c**.

We observed distinct contributions from each component in the GRASP framework. Removing heterogeneous graph features *GRASP*_*-HG*_ leads to a noticeable decline across nearly all metrics for both mRNA and lncRNA, underscoring the importance of substructure-aware graph representations and highlighting their central role within the overall model framework. Excluding sequence features leads to a moderate decrease in performance, demonstrating that k-mer and DACC features provide valuable nucleotide-level information that complements graph-based embeddings. Omitting the label correlation constraint *GRASP*_*-cc*_ generally leads to smaller performance degradation. However, the impact is more pronounced on the MCC metric, which drops from 0.302 to 0.248 for mRNA and from 0.234 to 0.185 for lncRNA, highlighting that label correlation primarily contributes to capturing complex multi-label dependencies rather than general predictive accuracy. The full GRASP model, which integrates heterogeneous graph and sequence features and label-correlation regularization, consistently achieves the highest scores across major indicators, confirming the synergistic effect of these components in enhancing prediction accuracy and reliability.

### 3.4 Global importance analysis reveals structural drivers of RNA localization

To better understand GRASP’s decision-making mechanism and investigate the biological relevance of the learned representations, we conduct a substructure-level importance analysis. In this analysis, we quantify the contribution of each node to the model’s predictions by computing substructure importance scores. Specifically, gradients of base, loop, and stem node features with respect to the target labels are backpropagated through the heterogeneous graph convolution network. Following the Grad × Input method (absolute value of the gradient multiplied by the activation, then summed) (32,33), we quantify the contribution of each substructure and base to the model’s predictions.

By performing a comprehensive substructure-level importance analysis across the entire lncRNA and mRNA datasets, we show the distribution of importance scores for the top 10 important loops and stems in each RNA sequence (**Figure 4a & b**). Among high-importance substructures, stems generally contribute more than loops, with right-skewed distributions, suggesting that a small subset of substructures may carry greater importance. Furthermore, **Figures 4c** and **e** summarize the relative importance across node types, for lncRNAs, stem loop base, whereas for mRNAs, stem importance dominates, but loops and bases are nearly equal. These results indicate that the model tends to assign higher importance to stem regions while still leveraging information from loops and bases, with minor RNA-type-specific differences. We then analyze importance variations among different loop subtypes (**Figure 4d** and **f**). Multi-loops generally show the highest importance, especially in lncRNAs. In mRNAs, hairpin loops are relatively more important than they are in lncRNAs, though multi-loops remain the most influential, with internal loops contributing least. This indicates that the model captures subtype-specific relevance patterns, which may reflect diverse functional roles. Finally, we examine substructure importance as a function of average base-pairing distance (**Figure 4g** and **h**). Importance generally declines with increasing distance, indicating a preference for short-range interactions, although some long-range substructures retain moderate importance. Notably, this trend is less pronounced in mRNAs, suggesting that the model captures long-range interactions more effectively in mRNAs, which may contribute to its superior performance on this dataset.

**Figure 4.**
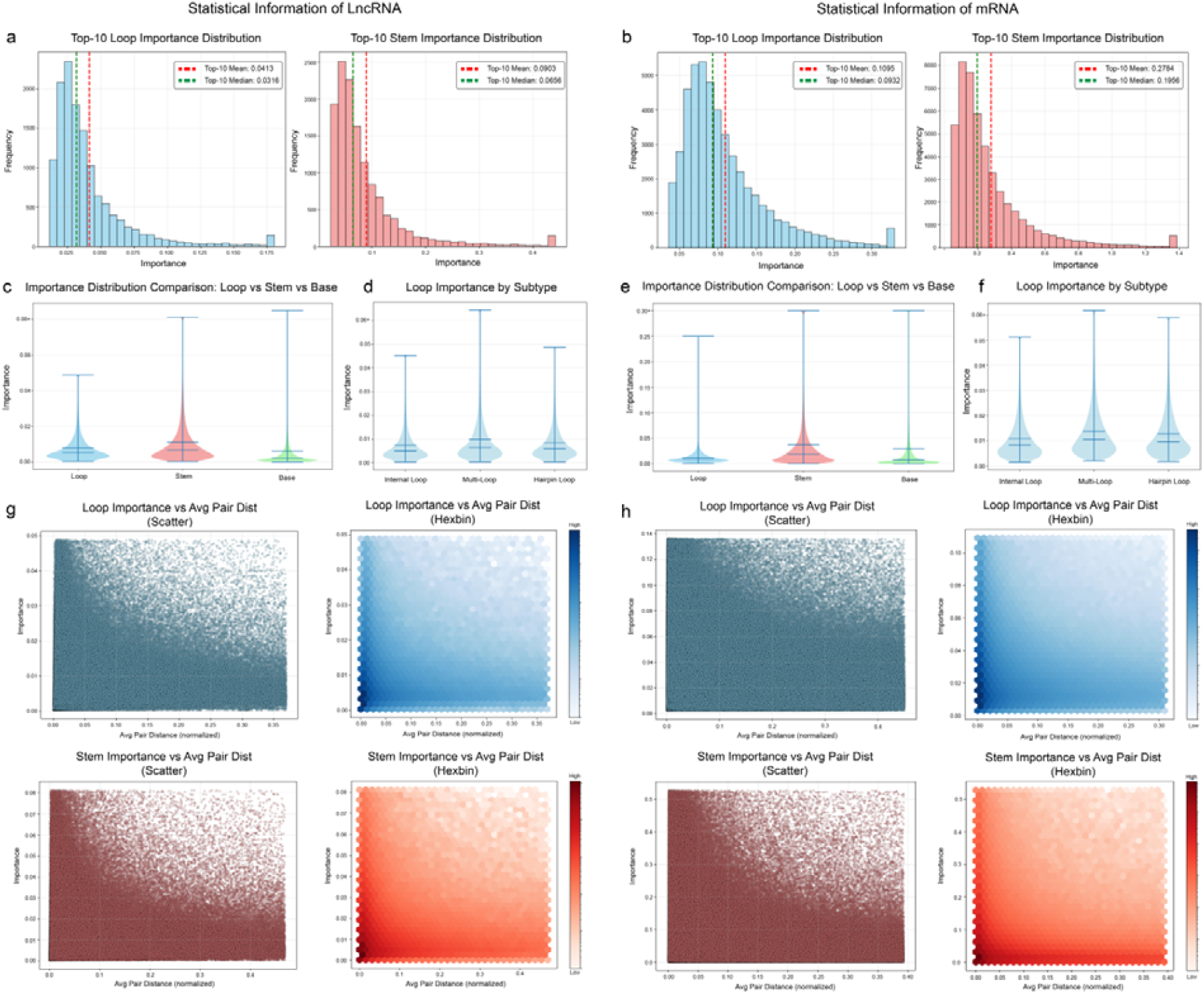
Dataset-level visualization of substructure importance on lncRNA and mRNA datasets. (a, b) Distribution of importance for the top10 important loops and stems in each RNA sequence, for lncRNA (a) and mRNA (b). (c, e) Comparison of importance scores among different node types for lncRNA (c) and mRNA (e). (d, f) Comparison of importance among loop subtypes on lncRNA (d) and mRNA (f). (g, h) Relationship between substructure importance and average base-pairing distance on lncRNA (g) and mRNA (h).

### 3.5 Important substructures are enriched for RNA modification sites

To assess whether the substructural patterns identified as important by GRASP correspond to biologically meaningful regions, we evaluate their enrichment in experimentally validated RNA modification sites. RNA modification annotations are retrieved from the RMBase 3.0 database (34). In total, the analysis includes 2279 lncRNA sequences with 32,135 annotated modification sites and 21,345 mRNA sequences with 610,124 annotated modification sites. For this analysis, sequences enriched for modification sites within the top 40% are reserved as the test set, while the remaining sequences are used for model training. For each test sequence, substructures are ranked by importance and divided into ten fixed segments using global index boundaries based on the average structure count. Coverage efficiency is then assessed by quantifying the overlap between modification sites and substructures in each segment.

We performed similar analyses on both lncRNA (**Figure 5**) and mRNA (**Figure 6**) datasets. Substructures are ranked by importance and divided into ten bins (**Figure 5a** and **Figure 6a**). In both RNA types, higher-importance substructures exhibit greater per-structure coverage, although the decline in coverage with decreasing importance is somewhat less pronounced for mRNAs. **Figure 5b** and **Figure 6b** show the relationship between substructure importance and marginal efficiency, which measures the additional coverage each substructure provides relative to its share of total structures. The results indicate that top-ranked, high-importance substructures cover many modification sites per structure, while lower-ranked ones contribute progressively less. **Figure 5c** and **Figure 6c** show one representative sequence each for lncRNA and mRNA, respectively, demonstrating both the global distribution of substructure importance and the nucleotide-level importance within selected high-importance substructures. Together, these results reveal a trend in both RNA types: substructures with higher importance scores tend to cover a larger fraction of modification sites, highlighting the model’s ability to prioritize biologically relevant regions and supporting the interpretability of the learned importance scores.

**Figure 5.**
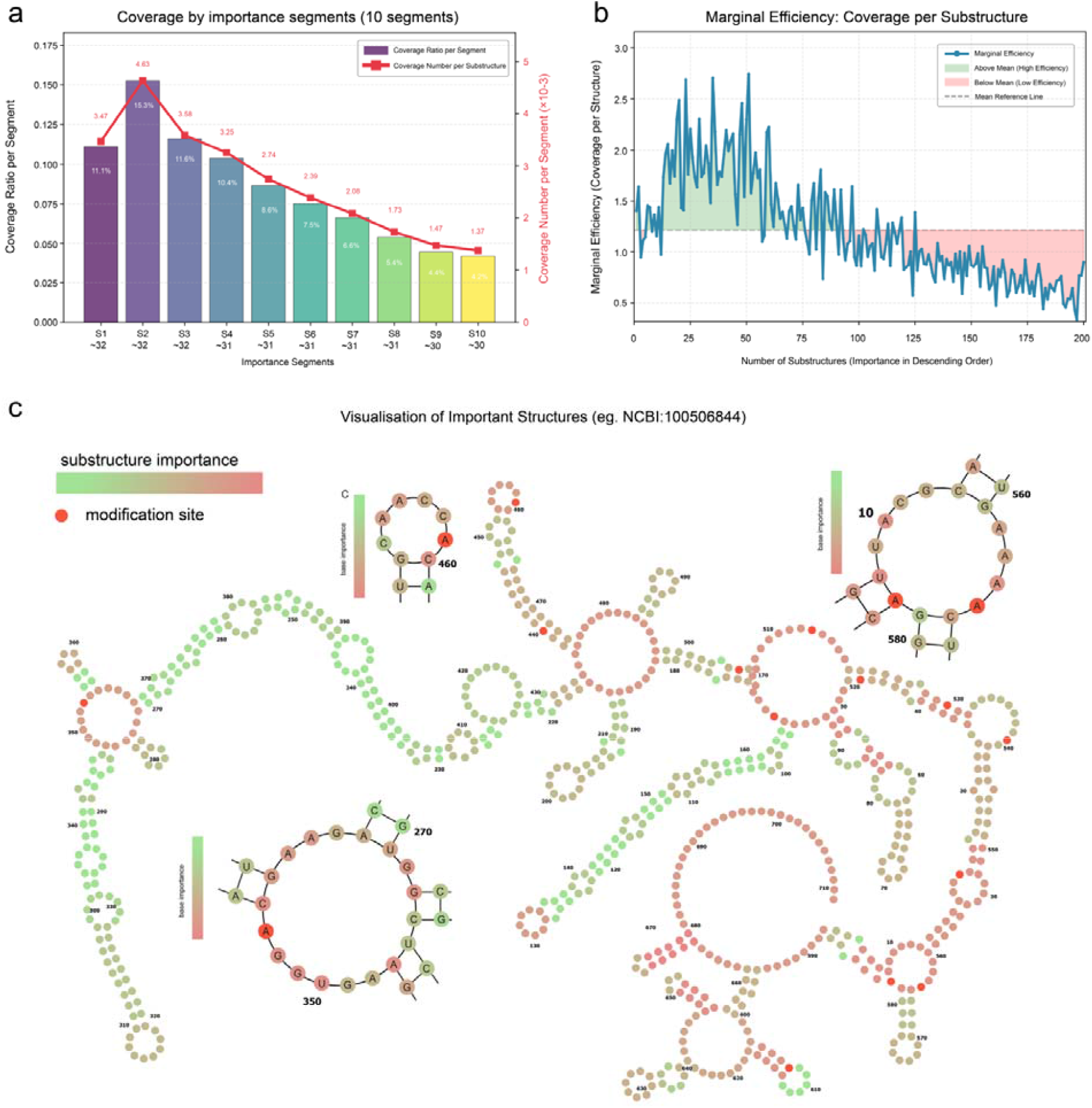
Biological relevance of identified important substructures on the lncRNA dataset. (a) Stratified coverage of RNA modification sites across ten importance-ranked substructure bins. (b) Marginal efficiency of individual substructures in covering modification sites. (c) Visualization of a representative lncRNA (NCBI:100506844), with global secondary structure colored by substructure importance and detailed base-level importance shown for selected substructures.

**Figure 6.**
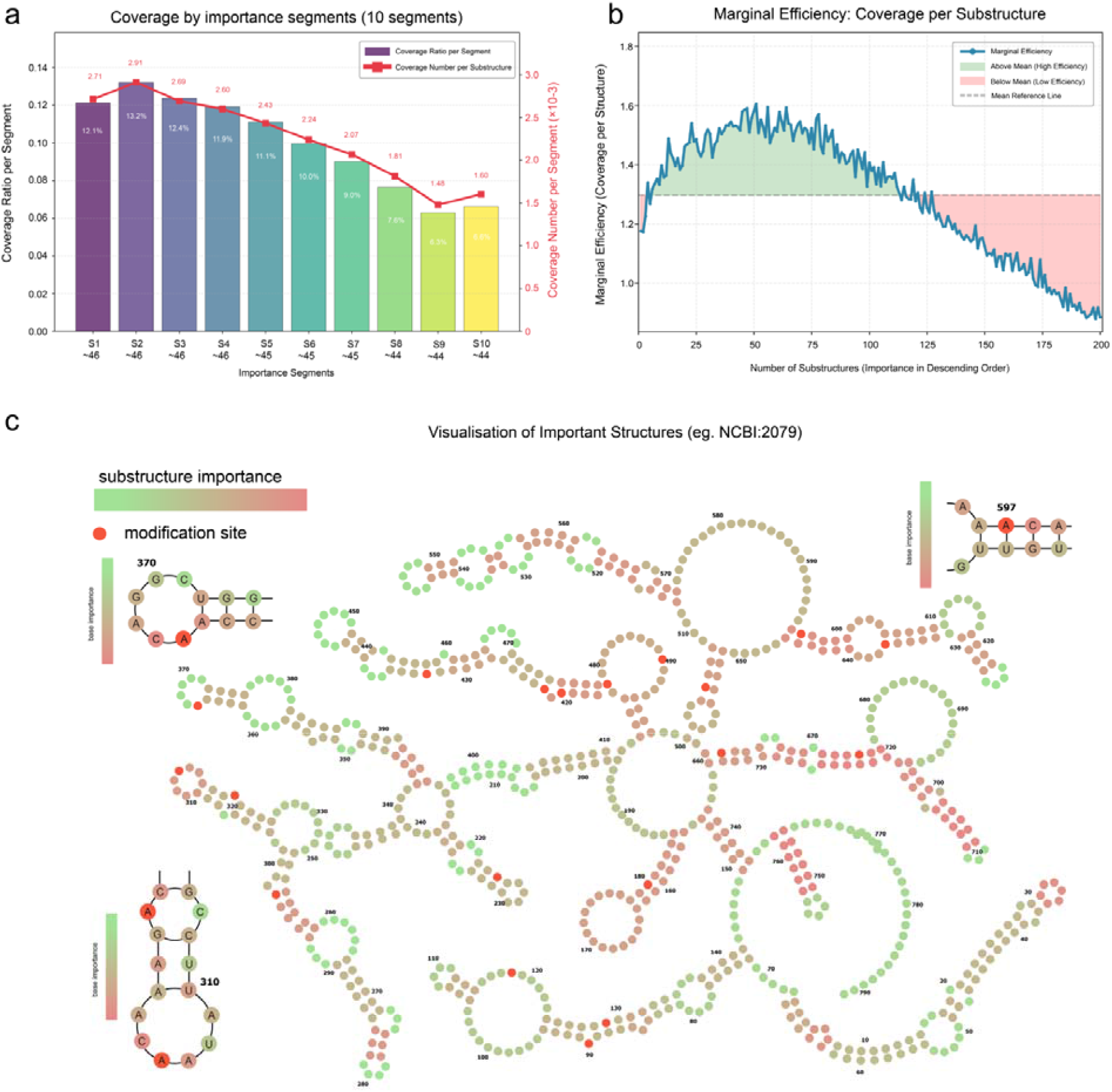
Biological relevance of identified important substructures on the mRNA dataset. (a) Stratified coverage of RNA modification sites across ten importance-ranked substructure bins. (b) Marginal efficiency of individual substructures in covering modification sites. (c) Visualization of a representative mRNA (NCBI:2079), with global secondary structure colored by substructure importance and detailed base-level importance shown for selected substructures.

### 3.6 Genome-wide compartment prediction and biological function analysis

To evaluate the broader applicability of GRASP, we perform genome-wide predictions on all human mRNA and lncRNA sequences. All transcript sequences are downloaded from GENCODE (35), with any transcripts corresponding to genes present in the benchmark datasets excluded, resulting in 50,128 mRNAs and 122,858 lncRNAs. GRASP is then applied to predict their subcellular localization probabilities across eight compartments. For mRNAs, a higher confidence threshold (0.9) is used due to generally stronger prediction reliability, whereas a slightly lower threshold (0.8) is applied for lncRNAs to ensure a sufficient number of candidate RNAs for downstream analysis given their overall lower prediction confidence scores, the distribution of predicted subcellular localization is illustrated in **Figure 7**. We next examine the functional relevance of these predicted localizations. For mRNAs, Gene Ontology (GO) enrichment analysis(36) is directly performed using Metascrape. For lncRNAs, due to the limited availability of direct functional annotations, AnnoLnc 2.0 (37) is used to infer potential functional associations via the GO annotations of their co-expressed genes. This analysis allows the identification of functional patterns corresponding to each predicted subcellular compartment. The resulting enrichment profiles are summarized in **Figure 8**, revealing that the predicted localizations correspond to biologically meaningful functional tendencies for both mRNAs and lncRNAs (2,38).

**Figure 7.**
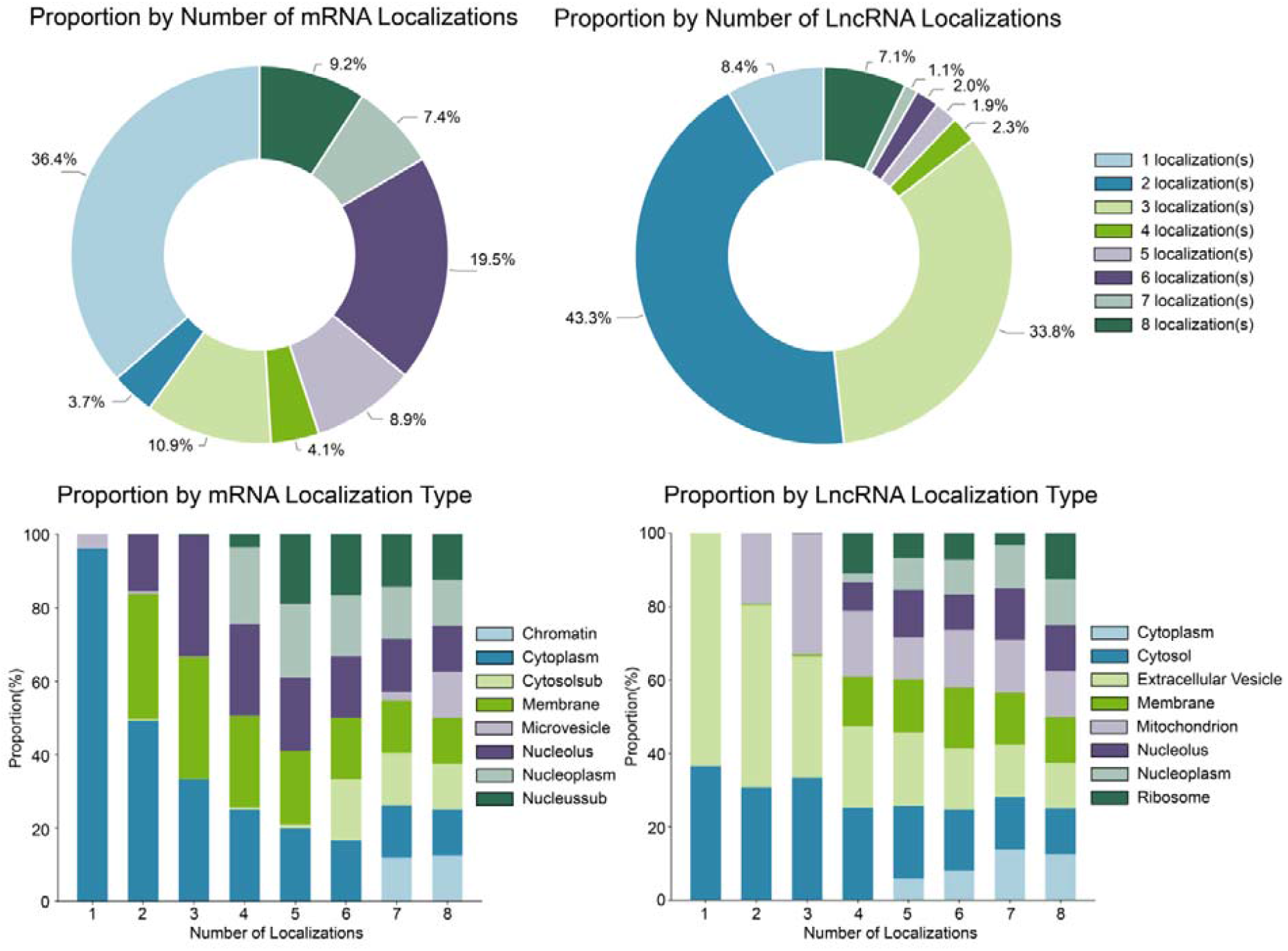
The pie chart shows the proportion of transcripts with different numbers of predicted subcellular localizations. The stacked bar chart further illustrates, for each localization count, the relative contribution of individual subcellular compartments, revealing how specific localization types are distributed among transcripts with different numbers of predicted localizations.

**Figure 8.**
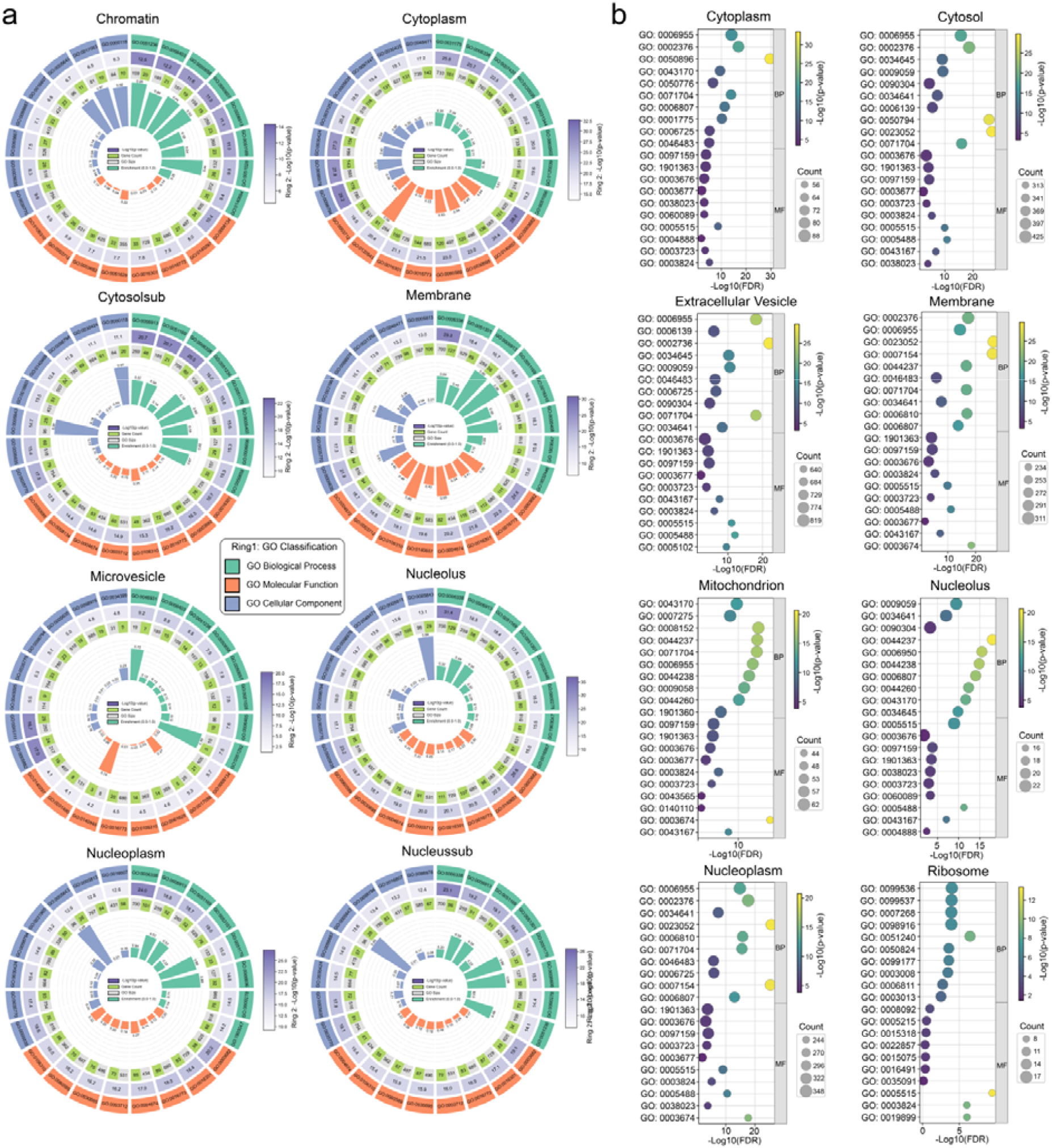
GO enrichment analysis results of predicted transcripts across subcellular localizations. (a) Results of mRNAs assigned to eight subcellular localizations based on direct GO enrichment analysis. (b) Results of lncRNAs assigned to eight subcellular localizations, with functions inferred via co-expressed genes.

**Figure 8a** illustrates GO enrichment of mRNAs. Chromatin-localized mRNAs are enriched for RNA transport (GO:0050658) and ribonucleoprotein granules (GO:0035770), reflecting co-transcriptional processing. Cytoplasmic mRNAs associate with neuron projection development (GO:0031175) and postsynaptic components (GO:0098794), supporting localized translation. Cytosolic mRNAs show strong links to nucleocytoplasmic transport (GO:0006913) and chromatin remodeling (GO:0006338). Membrane-associated mRNAs are connected to cell division (GO:0051301) and nuclear transport (GO:0051169). Microvesicle mRNAs relate to RNA localization (GO:0006403) and pore complex assembly (GO:0046931). Nuclear compartment mRNAs consistently exhibit nucleocytoplasmic transport (GO:0006913) and chromatin binding (GO:0003682). While most enrichments reflect the biological roles of each compartment (38-41), some associations, particularly the widespread enrichment of kinase activity (GO:0016301) in nuclear regions, may indicate functional translation sites rather than final protein activity locations (42).

**Figure 8b** illustrates GO enrichment of lncRNAs, cytoplasmic and cytosolic lncRNAs are strongly associated with immune response (GO:0006955) and protein/RNA binding (GO:0005515, GO:0003723). Extracellular vesicle lncRNAs show enrichment for cell-cell signaling (GO:0023052) alongside similar binding functions. Membrane-associated lncRNAs participate in signal transduction (GO:0007165) and immune system processes (GO:0002376). Mitochondrial lncRNAs are linked to metabolic processes (GO:0008152), while nucleolar lncRNAs specifically relate to RNA metabolism (GO:0016070) and biosynthesis (GO:0009059). Nucleoplasmic lncRNAs share features with both immune and metabolic regulation. Ribosome-localized lncRNAs are notably connected to synaptic signaling (GO:0099536). The pervasive enrichment of binding functions across all compartments accurately reflects lncRNAs’ core molecular mechanisms, while specific process enrichments validate the functional relevance of their predicted subcellular distributions(43-45).

## 4. Conclusion and future perspectives

While numerous models have been proposed to predict mRNA and lncRNA subcellular localization, most of these approaches do not deeply specialize in modeling RNA secondary structures and are typically limited to a single RNA type. In this study, we present GRASP, a novel graph-based representation model designed for RNA subcellular localization prediction. By converting RNA sequences into heterogeneous graphs, our approach enables the fusion of base-level and substructure-level information, such as loops and stems, to learn comprehensive RNA representations. This model integrates original sequence data with structural features, offering a more holistic understanding of RNA localization. Furthermore, we introduced the ASL loss function with a label-dependent regularization term to improve model performance by capturing co-occurrence patterns among subcellular localization sites. Compared with state-of-the-art specialized models and widely used RNA foundation models, GRASP achieves superior performance in RNA subcellular localization prediction, particularly in micro recall, indicating its enhanced ability to identify more true-positive instances. Importantly, this improvement is achieved while maintaining competitive precision and concurrently increasing Average MCC, reflecting not only better overall predictive performance but also robust handling of the data imbalance inherent in multi-label prediction tasks. The high MCC further indicates the model’s effectiveness in capturing complex label dependencies, highlighting its suitability for multi-label RNA subcellular localization prediction. Through extensive ablation studies, we demonstrate that the heterogeneous graph representation strategy enhances the model’s feature-learning capabilities, enabling more accurate and interpretable predictions. Nevertheless, sequence information remains a core component, consistent with our previous findings (17). This highlights the complementary nature of sequence and structure in RNA function prediction. Additionally, our interpretability analyses offer deeper insights into the model’s decision-making process. Notably, in both lncRNA and mRNA, the model consistently captures strong predictive signals from stem regions, suggesting that these paired structures play a central role in determining subcellular localization. These analyses also confirm the model’s capability to utilize both short-range and long-range base pairings, even when short-range interactions are dominant, highlighting its sensitivity to hierarchical structural features critical for RNA function. The modification site coverage analysis further validates GRASP’s ability to identify biologically relevant substructures. In large-scale human genome-wide predictions of mRNA and lncRNA subcellular localization, along with Gene Ontology (GO) enrichment analysis, the predicted substructures showed strong alignment with known functional enrichments across cellular compartments, further validating the biological relevance of the model’s predictions. The results also highlight functional distinctions between mRNAs and lncRNAs: mRNAs are primarily involved in RNA transport, nucleocytoplasmic trafficking, and localized translation, whereas lncRNAs exhibit broader regulatory and signaling roles, including immune responses, metabolic regulation, and pervasive binding functions. Note that the lncRNA functions described here are inferred from co-expression with annotated genes, and more precise downstream interpretations will require expanded GO annotations. In summary, GRASP is a powerful and practical tool for predicting RNA subcellular localization, and it can also facilitate downstream studies, such as RNA modification analysis and functional characterization.

However, it is important to note that the current model does not incorporate tertiary RNA structure due to computational and efficiency constraints(46,47). We plan to explore this in future work to further enhance the model’s ability to capture higher-order structural features. While previous specialized models have provided invaluable insights, there is no doubt that RNA research will benefit from more generalized models, including those leveraging language model architectures. We will continue to focus on the development of universal models that can expand the scope of downstream tasks, making them applicable to a wider range of RNA research.

## Data availability

The GRASP web server is publicly available at http://grasp.biotools.bio. The source code and the data underlying this article are publicly available on GitHub (https://github.com/ABILiLab/GRASP).

## Supplementary data

Supplementary Data are available at NAR Online.

## Funding

This work is supported by the National Key Research and Development Program of China (No. 2022YFF1000100), the National Natural Science Foundation of China (No. 62202388), and the Qin Chuangyuan Innovation and Entrepreneurship Talent Project (No. QCYRCXM-2022–230). F.L. is supported by the Australia National Health and Medical Research Council (NHMRC) Investigator Fellowship (2041439).

## Conflict of interest statement

None declared.

## Notes

https://github.com/ABILiLab/GRASP

### Competing Interest Statement

The authors have declared no competing interest.

