## Supplementary materials for "Graph-based RNA structural representation reveals determinants of subcellular localization"

**1. Evaluation metrics**

The example-based metrics include example accuracy(${ACC}_{example}$), average precision score($Average Precision$), hamming loss($Hamming Loss$), ranking loss($Ranking Loss$), zero-one loss($One-error$), and coverage error($Coverage$), which are defined as:

$${ACC}_{example}=\frac{1}{N}\sum_{i=1}^{N} \left| \left. \frac{Y_{i}\cap P_{i}}{Y_{i}\cup P_{i}} \right| \right.$$

(1)

$$Average Precision=\frac{1}{L}\sum_{j=1}^{L} \sum_{k=1}^{N} P_{j,k}\left( R_{j,k}-R_{j,k-1} \right)$$

(2)

$$Hamming Loss=\frac{1}{N\cdot L}\sum_{i=1}^{N} \sum_{j=1}^{L} 1\left( y_{i}\left[ j \right]\neq\hat{y_{i}}\left[ j \right] \right)$$

(3)

$$Ranking Loss=\frac{1}{N}\sum_{i=1}^{N} \frac{1}{\left| P_{i} \right|\cdot\left| \bar{P}_{i} \right|}\sum_{j\in P_{i}} \sum_{k\in\bar{P}_{i}} 1\left( f_{i} \left( j \right)\leq f_{i} \left( k \right) \right)$$

(4)

$$One-error=\frac{1}{N}\sum_{i=1}^{N} 1\left( \underset{j\in\left\{ 1,2\ldots,L \right\}}{argmax} \left( f_{i} \left( j \right) \right)\notin P_{i} \right)$$

(5)

$$Coverage=\frac{1}{N}\sum_{i=1}^{N} \left( \max_{j\in Y_{i}} \left( {rank}_{i}\left( j \right) \right)-1 \right)$$

(6)

where $N$ is the number of samples and $L$ is the number of labels, $Y_{i}$ and $P_{i}$ denote the true label set and the predicted label set of $i$-th label, respectively, $y_{i}\left[ j \right]$ and $\hat{y_{i}}\left[ j \right]$ represent the true and predicted value of the $i$-th sample in $j$-th label, $f_{i} \left( j \right)$ denotes the predicted score corresponding to the $i$-th sample in $j$-th label, ${rank}_{i}\left( j \right)$ denots the rank of $j$-th label in the order of predicted scores of the $i$-th sample, $P_{j,k}$ and $R_{j,k}$ represent the precision and recall of the first $k$ samples on the $j$-th label.

The label-based metrics include average F1-score($AvgF1$), micro precision($Micro Precision$), micro recall($Micro Recall$), average AUC($AvgAUC$), and average matthews correlation coefficient($AvgMCC$), which are defined as:

$$AvgF1=\frac{1}{L}\sum_{l=1}^{L} \frac{2\cdot{TP}_{l}}{2\cdot{TP}_{l}+{FP}_{l}+{FN}_{l}}$$

(7)

$$Micro Precision=\frac{\sum_{l=1}^{L} {TP}_{l}}{\sum_{l=1}^{L} \left( {TP}_{l}+{FP}_{l} \right)}$$

(8)

$$Micro Recall=\frac{\sum_{l=1}^{L} {TP}_{l}}{\sum_{l=1}^{L} \left( {TP}_{l}+{FN}_{l} \right)}$$

(9)

$$AvgAUC=\frac{1}{L}\sum_{l=1}^{L} \frac{\sum_{i\in P_{l}} \sum_{j\in N_{l}} 1\left( f_{i} \left( l \right)>f_{j} \left( l \right) \right)+0.5\cdot1\left( f_{i} \left( l \right)=f_{j} \left( l \right) \right)}{\left| P_{l} \right|\cdot\left| N_{l} \right|}$$

(10)

$$AvgMCC=\frac{1}{L}\sum_{l=1}^{L} \frac{{TP}_{l}\cdot{TN}_{l}-{FP}_{l}\cdot{FN}_{l}}{\sqrt{\left( {TP}_{l}+{FP}_{l} \right)\left( {TP}_{l}+{FN}_{l} \right)\left( {TN}_{l}+{FP}_{l} \right)\left( {TN}_{l}+{FN}_{l} \right)+\varepsilon}}$$

(11)

where $P_{l}$ and $N_{l}$ denote the sets of positive and negative samples for the $l$-th label, respectively, and $\left| \cdot\right|$ represents the cardinality of a set. ${TP}_{l}$, ${FP}_{l}$, ${TN}_{l}$, and ${FN}_{l}$ refer to the numbers of true positives, false positives, true negatives, and false negatives for the $l$-th label, respectively, $\varepsilon$ is a smoothing term to prevent the denominator from being zero.
